# Adaptive laboratory evolution of a minimal cell to low temperature

**DOI:** 10.64898/2026.06.03.730023

**Authors:** Masaki Mizutani, Minoru Moriyama, Ryuichi Koga, Takema Fukatsu, Shigeyuki Kakizawa

## Abstract

Temperature is a fundamental constraint on life. Most organisms adapt to temperature change through complex regulatory and stress-response systems. Whether such adaptation is possible in a genomically minimal cell remains unclear. Here, adaptive laboratory evolution was performed on the minimal genome bacterium JCVI-syn3B, shifting growth from 37°C to 25°C, a temperature at which the original strain cannot grow. After 40 serial passages, the evolved strains exhibited robust growth at 25°C. Multi-omics analyses revealed the adaptation mechanisms including increased mRNA turnover, enhanced DNA unwinding and stabilization of replication intermediates, elevated glycerolipid synthesis, and upregulation of division proteins associated with Z-ring assembly. Whole-genome transplantation was performed to distinguish genetic or non-genetic contributions, demonstrating that the cold-adapted phenotype was largely genetically encoded. These results indicated that even a minimal cell retains substantial evolvability and highlight its potential as a tractable research platform for cellular and evolutionary biology.

## Main

Temperature is a critical determinant for survival of all organisms. Each organism is adapted to an optimal growth temperature characteristic of its own habitat, and sudden shifts in temperature are generally linked to a diminished probability of survival. Various mechanisms have been identified that enable organisms to respond and adapt to changes in growth temperature, including active heat generation in homeotherms ^1,2^, reduced energy consumption through hibernation ^3,4^, maintenance of membrane fluidity through changes in fatty acid composition ^5^, maintenance of protein flexibility through amino acid substitution ^6^, and the preservation of protein function via the expression of heat shock proteins, cold shock proteins, and molecular chaperones ^7–10^. Elucidation of these mechanisms had contributed to understanding survival strategies and fundamental cellular functions of organisms.

Whether environmental adaptation is possible in a cell lacking most stress-response and regulatory systems remains unclear. The genome-streamlined bacterium JCVI-syn3B ^11–15^ provides a tractable model to uncover adaptive mechanisms that are normally masked by complex regulatory networks. Here, we show that JCVI-syn3B cells rapidly adapted to low temperature by changing cellular processes derived from only several mutations in genome. We performed adaptive laboratory evolution to reoptimize the growth temperature from 37 °C to 25 °C, a temperature at which the original strain cannot grow. Multi-omics analyses revealed four major processes reorganized to low temperature adaptation; RNA degradation, DNA replication, glycerolipid synthesis, and cell division. The genome transplantation analysis demonstrated that these reorganizations of cellular processes were based on the mutations of the genome. These findings showed the evolvability of genome-streamlined bacterium to non-optimal environments, and the potential of availability for understanding fundamental mechanisms of cellular adaptation and evolution across different environments.

### Growth speed of minimal cell at various temperatures

JCVI-syn3B cells were cultured at 37°C, 33°C, 30°C, and 27°C in SP4 medium, which were monitored for growth by the absorbance at 560 nm for six days. The absorbance value decreased by the accumulation of acidic metabolites from glycolysis, reflecting the growth of cultures ^16^. The growth curve reached a plateau after approximately two and four days at 37°C and 33°C, respectively. Growth at 30°C was substantially slower and did not reach a plateau within six days, and no detectable growth was observed at 27°C (Fig. 1A). The doubling times at 37°C, 33°C, and 30°C were 2.6 ± 0.1, 6.2 ± 0.5, and 15.9 ± 1.1 hours, respectively (Fig. 1B).

**Figure 1.**
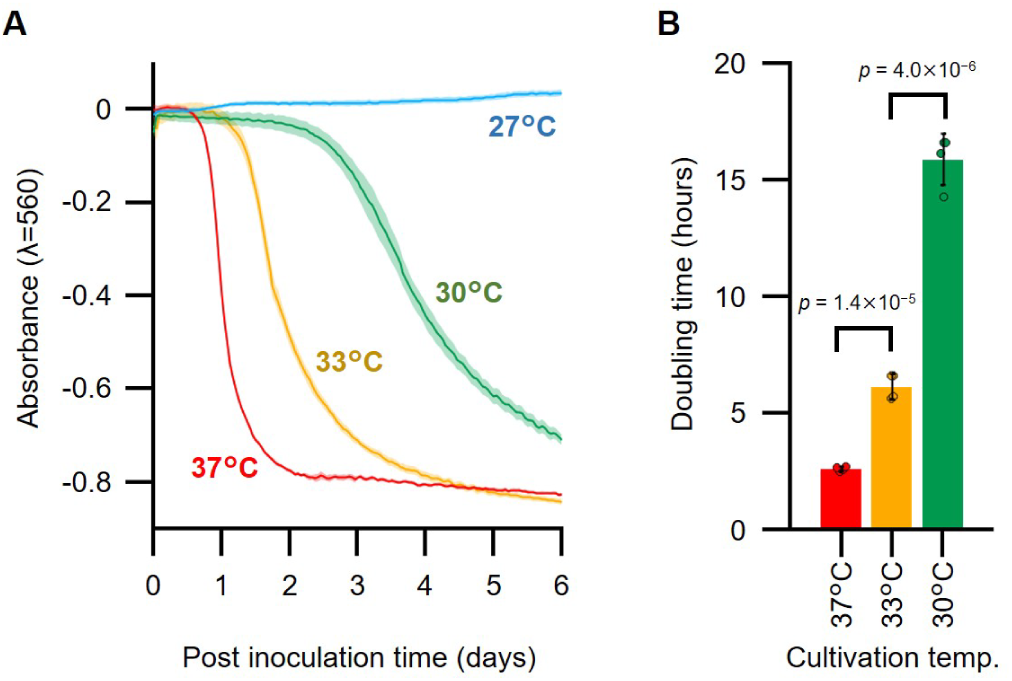
Growth speed of original JCVI-syn3B strain at various temperatures. (A) Growth curve. The absorbance values at 560 nm of cultures incubated at 37°C, 33°C, 30°C, and 27°C were measured with one-hour intervals and the average values of four cultures are shown as curved lines in red, yellow, green, and cyan, respectively. Light-coloured lines indicate the standard deviations. (B) Doubling time in the early exponential growth phase at each temperature. Individual values and standard deviations are shown as filled circles and error bars, respectively. *P* values of Student’s *t*-test are upper of graph.

### Adaptive laboratory evolution to low temperature

To investigate the evolvability of JCVI-syn3B to adapt non-optimal environments, cells were subjected to serial passaging at 30°C followed by 25°C in four parallel lineages. The JCVI-syn3B was first cultured at 37°C in SP4 medium, then a part of grown culture was transferred to fresh SP4 medium and incubated at 30°C until late-exponential or stationary phase (passage 1). Then, a part of grown culture was again transferred to fresh SP4 medium and incubated at 30°C (passage 2). This transfer was repeated up to passage 20 in four parallel lineages (30-L1 to 30-L4). The cultivation time of passage 1 and 2 averaged 70 hours across four lineages. Passages 3–5 required approximately 105 hours, longer than passage 2, suggesting that the passage 1 and 2 strains carried over the high growth activity presumably derived from the parental strain cultured at 37°C. Passages 6–17 required 58–76 hours, and passages 18–20 required about 46 hours (Fig. 2). These results indicate that JCVI-syn3B cells adapted their growth to low temperature through continuous passaging. For further investigation, each lineage of passage 20 at 30°C was transferred to fresh SP4 medium and incubated at 25°C (passage 1 at 25°C). Even though the original strain could not grow at 25°C, the passaged strains could grow. The passaging process was repeated again at 25°C up to passage 20 in four parallel lineages (25-L1 to 25-L4). From passages 2–12, the cultivation time was mostly over 200 hours, with an average of 250 hours, whereas from passages 13–20, it was mostly under 155 hours, with an average of 149 hours (Fig. 2). These results indicate that JCVI-syn3B can beyond the limitation of culture condition through adaptive evolution within only 40 serial passages.

**Figure 2.**
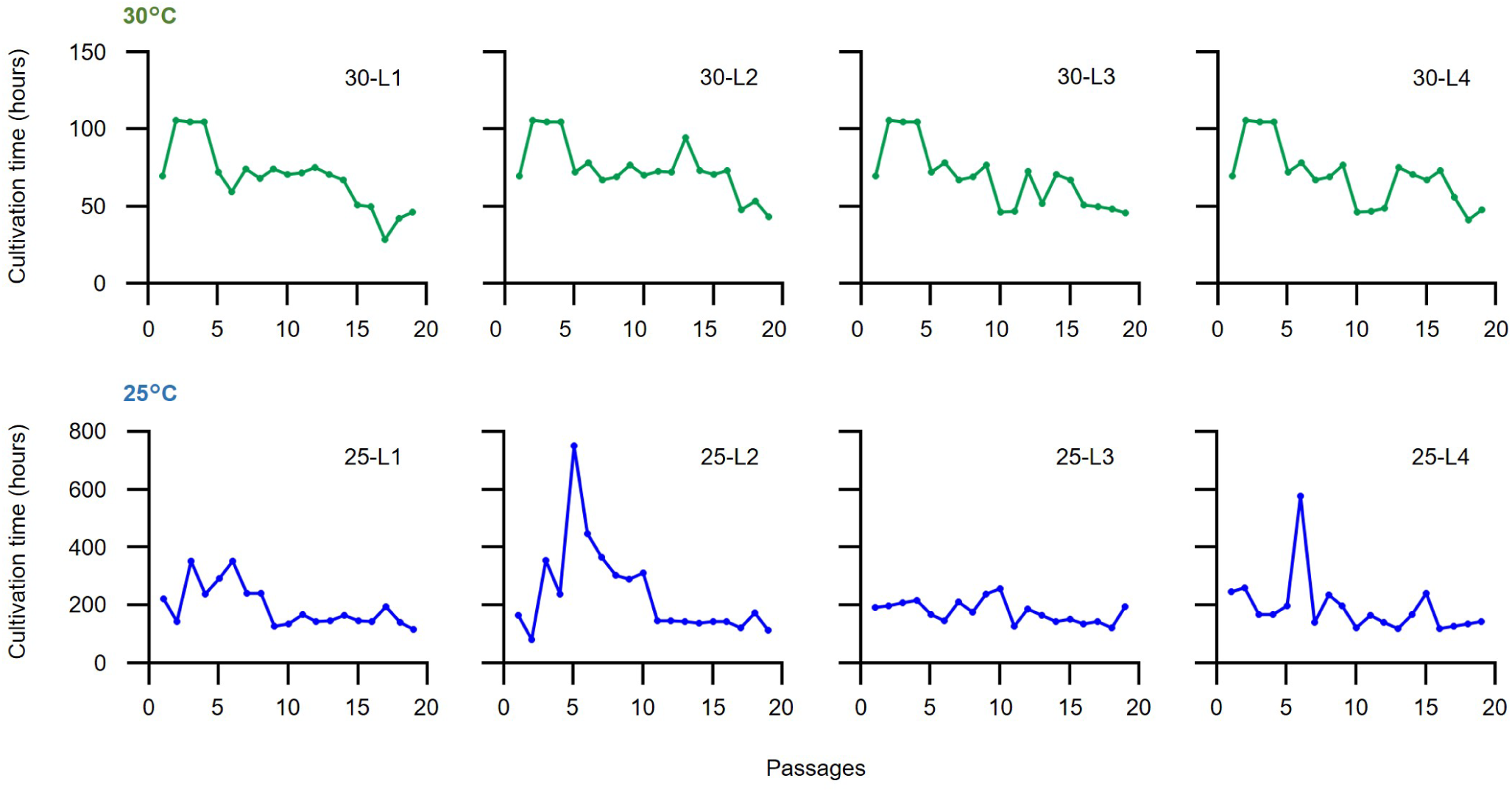
Changes in growth rates during adaptive laboratory evolution. The cultivation time required for each passage are plotted for the four independent lineages at 30°C (green) and 25°C (blue).

### Trade-off relationships of growth during temperature adaptation

To investigate the growth speed of the evolved strains, four 30°C-adapted strains and four 25°C-adapted strains were cultured at 37°C, 30°C, and 25°C (Extended Data Fig. 1). The doubling time of the original strain was 2.5 ± 0.2 hours at 37°C (Fig 3A). The growth of evolved strains at 37°C was clearly divided into two groups. Strains 30-L1, 30-L2, 25-L1, and 25-L3 showed similar doubling times to the original strain, at 88%, 107%, 115%, and 87%, respectively. In contrast, the doubling times of strains 30-L3, 30-L4, 25-L2, and 25-L4 were approximately twice as long as that of the original strain, at 200%, 219%, 202%, and 229%, respectively. At 30°C, all evolved strains grew about twice as fast as the original strain (Fig 3B). Only the 25°C-adapted strains could grow well within 14 days at 25°C (Fig 3C). All 30°C-adapted strains could not grow at 25°C, possibly because they had been stored as frozen stocks and required more than 14 days for regrowth. The 25-L4 strain showed the fastest growth among the four 25°C-adapted strains at 25°C. These results indicate that trade off of growth temperature was observed in several strains but the characters were not uniform as previously reported in the temperature adaptation of *E. coli* ^10^.

**Figure 3.**
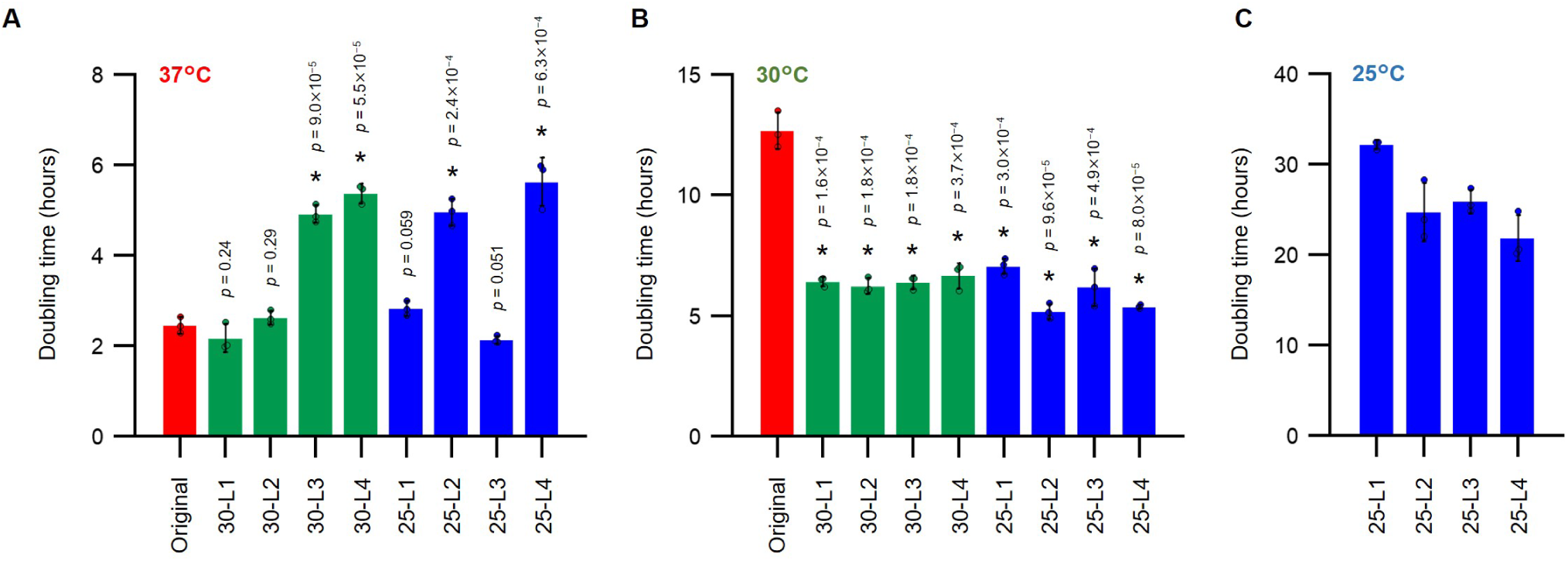
Doubling times of adapted strains at 37°C (A), 30°C (B), and 25°C (C). Individual values and standard deviations are shown as filled circles and error bars, respectively. *P* values of Student’s *t*-test against original strain are upper of graphs.

### Genome mutations for low-temperature adaptation

To determine the gene mutations associated with low-temperature adaptation, the genomes of original and four 25°C-adapted strains (25-L1 to 25-L4) were sequenced and detected mutations were summarized (Extended Data Table 1). Only 9–14 mutations were detected in each genome. Multiple mutations were identified across the four lineages in the RNA polymerase subunit beta (JOY36_02225), beta’ (JOY36_02220), and sigma factor (JOY36_01085) genes, ABC transporter genes (JOY36_00110, JOY36_00100, and JOY36_01045), suggesting that modifications of global transcription levels and uptake capabilities of certain substrates might be effective to cold adaptation. Furthermore, mutations were observed in three lineages within a functional unknown gene (JOY36_01130), implying a certain role of this gene under low-temperature conditions. Although several genes potentially important for cold adaptation were identified, the mechanism of cold adaptation could not be directly explained based solely on the observed genomic mutations.

### Transcriptome analysis

To examine transcriptional change after cold adaptation, RNA-seq analyses were performed for the original and four 25°C-adapted strains. In total 465 genes except transfer RNAs (tRNAs) and ribosomal RNAs (rRNAs) were analyzed. The principal component analysis for transcriptions per million (TPM) values indicated that low-temperature adaptation strategy had not only a particular tendency but also variations (Fig. 4). To simplify the transcription comparison, genes were classified into 31 functional categories (Extended Data Table 2 and Supplementary Information Table 1). Genes with high transcription levels in each strain were summarized (Extended Data Table 3). Overall, the two non-coding RNA (ncRNA) genes, transfer-messenger RNA (tmRNA) and the RNA component of RNase P, were highly expressed. In the original strain, these two genes alone accounted for 48.1% of the total RNA excluding tRNA and rRNA (Supplementary Information Table 1). Similar levels were observed in cold-adapted strains 25-L1, 25-L3, and 25-L4 (42.6%–51.7%), whereas strain 25-L2 showed a marked reduction (14.5%). The tmRNA (encoded by the *ssrA* gene) is essential for protein quality control and cellular homeostasis ^17^, whereas ribonucleases (RNase) P is involved in tRNA maturation ^18^, a process essential for protein synthesis and conserved across all domains of life.

**Figure 4.**
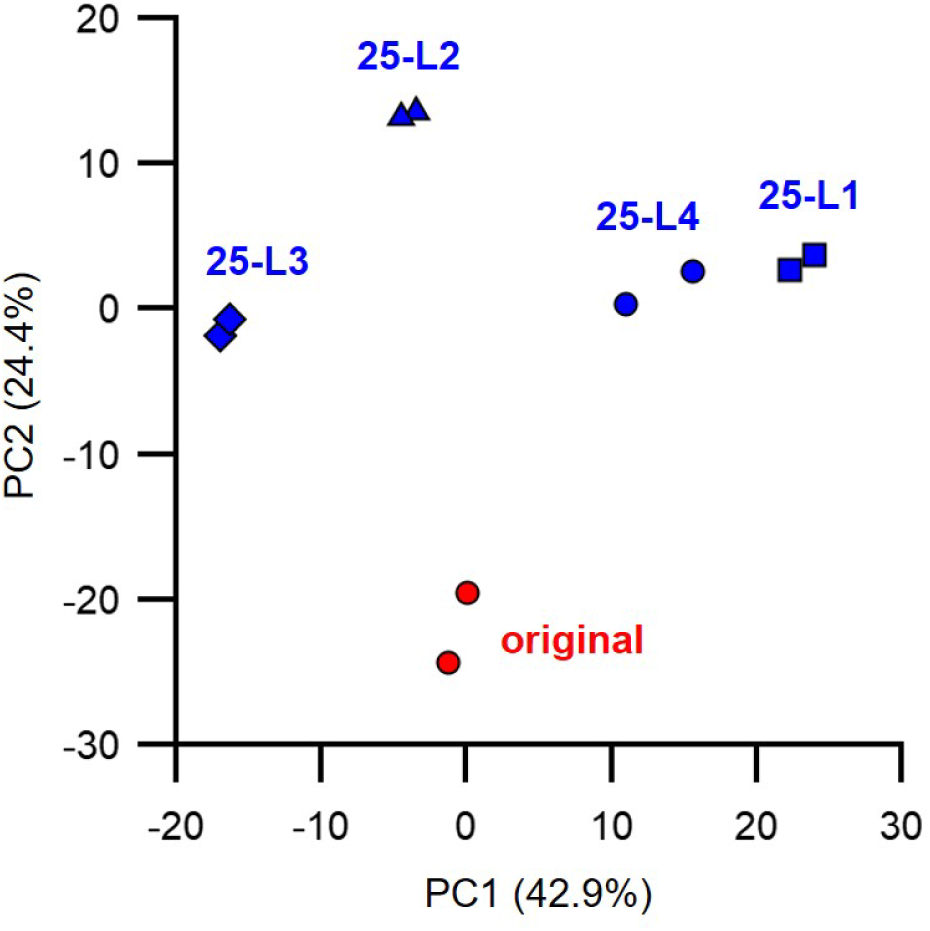
The principal component analysis of the RNA-seq analysis. Red circle: Original, Blue square: 25-L1, Blue triangle: 25-L2, Blue diamond: 25-L3, Blue circle: 25-L4. Two data set per strain.

The category showing the most substantial change in the cold-adapted strains was the phosphotransferase system (PTS), with an average expression level of 450% relative to the original strain. Since glycolysis is responsible for ATP production from glucose, this result suggests that ATP production capacity may have been enhanced in the cold-adapted strains. RNA degradation category was also highly expressed, with an average expression level of 365%. RNases have been proposed as key factors in the cold shock response of *E. coli*, functioning to degrade abnormally structured mRNAs and thereby facilitate efficient translation at low temperature ^7^. The F-type ATPase category showed an average expression level of 263%. Actually, mycoplasmas use F-type ATPase for not ATP synthesis but establishment of membrane potential. The lipid biosynthetic pathway from glycerol was generally overexpressed, with a particularly marked upregulation of the *glpK* gene, suggesting enhanced lipid metabolism during cold adaptation. These results suggest the involvement of multiple cold-adaptation mechanisms, including enhanced glycolysis, RNA degradation, and membrane potential.

### DNA methylation analysis

Whole-genome methylation analysis was performed for the original and two 25°C- adapted strains (25-L1 and 25-L4), and the representative bacterial methylations m4C and m6A were mapped to each genome (Supplementary Information Table 2). As a result, no genes showed a significant change in the number of methylation sites in either of the cold-adapted lineages compared with the original strain (Extended Data Table 4). Overall, both cold-adapted lineages showed a slight decrease in the total number of methylation sites, with the numbers of m4C and m6A being reduced by 3%–5% compared with the original strain. These results suggested that DNA methylation had a limited contribution in this cold adaptation.

### Proteomics analysis

To investigate the detailed mechanisms of cold adaptation, whole-cell proteomic analyses were performed using mass spectrometry. In total, 371 proteins (82% of the 455 coding proteins in the genome) were detected (Supplementary Information Table 3). To simplify the comparison of expression levels, proteins were classified into 31 functional categories, and the expression levels of each category are summarized (Extended Data Table 5). Proteins with high abundance in each lineage were summarized (Extended Data Table 6 and 7), and a certain degree of commonality was observed across lineages. The category with the highest expression ratio was cell division, with an average protein abundance of 307% compared to the original strain. In this category, four proteins, FtsZ, FtsA, SepF, and DivIVA were detected. Notably, the expression ratios of FtsZ, which forms a contractile ring and drives cell division through GTP hydrolysis, and SepF, which anchors FtsZ filaments to the cell membrane, ranged from 282% to 503% and 140% to 173% of the original strain, respectively. GTP hydrolysis activity decreases at low temperature, resulting in suppressed ring contraction. Therefore, the high expression of FtsZ and SepF likely support effective cell division under cold conditions. The second highest expression category was RNA degradation, consisting of six RNases. Notably, the expression ratios of the 3’-5’ exoribonuclease ranged from 316% to 413% of the original strain, and of RNase Y, RNase R, and two RNase J enzymes were also elevated ranged from 122% to 230%, consistent with the results of transcriptome. The two RNA biogenesis categories showed increased expression levels from 122% and 145%, corresponding to active RNA degradation and recycling. The expression ratios of glycerol kinase, which catalyzes the phosphorylation of glycerol to glycerol-3-phosphate for glycerol uptake, and transporters of cofactor, macromolecules, and ions were also upregulated, supporting more efficient nutrients transport at low temperature.

Focusing on other specific proteins, DNA helicase (DnaB) and single-stranded DNA-binding protein (SSB) showed high expression levels, ranging from 306% to 855% for DnaB and from 189% to 316% for SSB in strains 25-L2, 25-L3, and 25-L4. DnaB unwinds double-stranded DNA, while SSB stabilizes the resulting single strands, facilitating effective DNA replication under cold conditions, where DNA strand pairing is reinforced. Additionally, the expression level of the helicase-like enzyme UvrB increased in all lineages, ranging from 182% to 258%.

To examine the correlation between the RNA-seq and proteomics data, the expression levels from both datasets were plotted (Extended Data Fig. 2 and Supplementary Information Table 4). Overall, the expression patterns showed a modest correlation between the two analyses.

### Genome exchanges

Although only limited genomic differences were detected among the 25°C-adapted strains, proteomic and transcriptomic analyses revealed substantial changes, suggesting a potential contribution of cytoplasmic effects to cold adaptation. To test whether genomic mutations alone were sufficient, we performed genome transplantation ^19^. First, a puromycin resistance gene was inserted into the genomes of the original strain and the 25°C-adapted strain (25-L4) using the Cre/*lox* system ^16,20^. The genomic DNA from the both strains were then extracted and transplanted into the original cells. After transplantation, only the cells containing the 25°C-adapted genome formed colonies at 25°C (Fig. 5A). Next, the growth speed of the original, adapted, and transplanted strains, all of which carried the puromycin resistance gene, was compared at 25°C. The original strain showed no significant growth for 12 days (Fig. 5B). In contrast, both the 25-L4 and the transplanted strains exhibited clear growth, with similar doubling times of 20.3 ± 1.2 and 21.7 ± 1.3 hours, respectively (Fig. 5C). These results demonstrate that the genomic mutation was the crucial determinant for cold adaptation.

**Figure 5.**
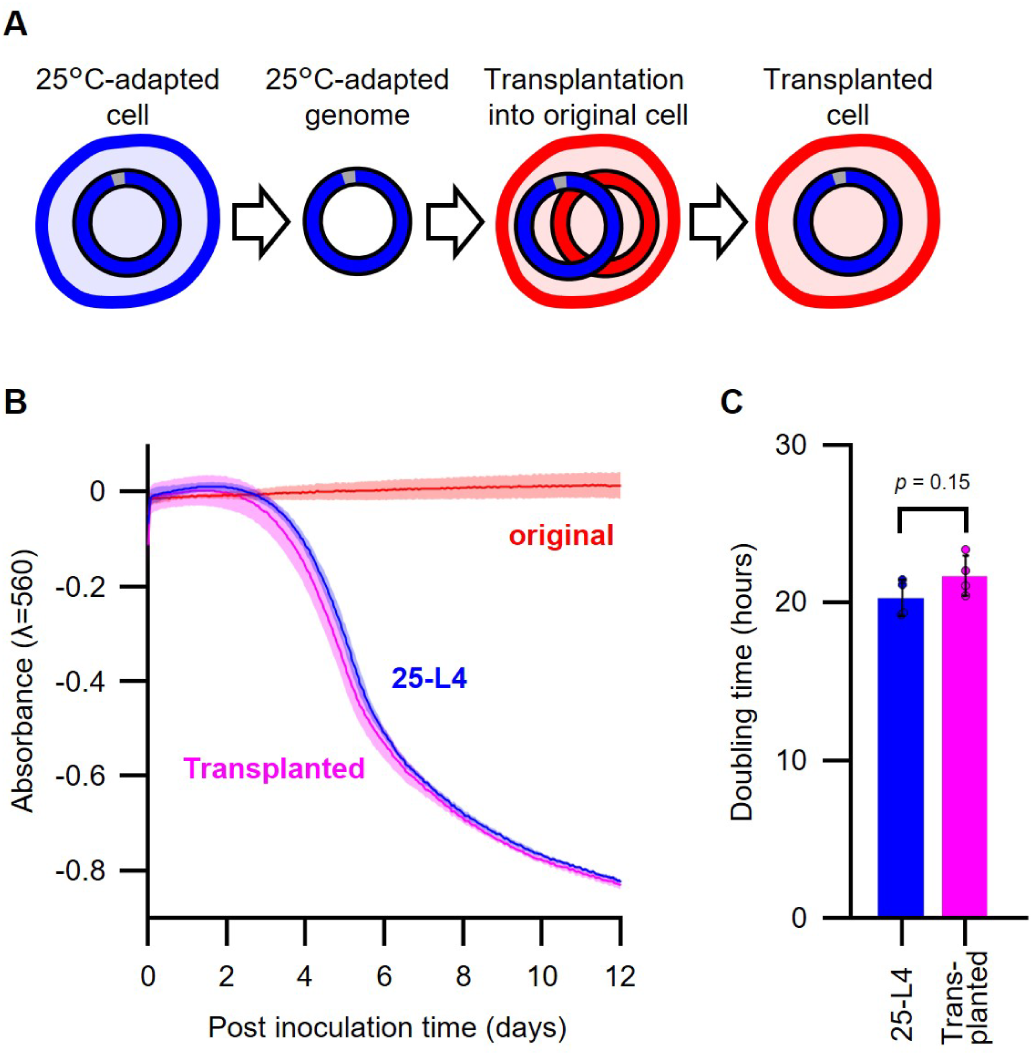
Whole genome transplantation experiments. (A) Summarized illustrations of genome transplantation. The puromycin-resistance gene was inserted into the genome, and transplanted cells were selected with puromycin. (B) Growth curves at 25°C. Mean values of four independent cultures are shown in red, blue, and magenta lines for the original, 25-L4, and genome-transplanted strains, respectively. Light-coloured lines indicate standard deviations. (C) Doubling times of 25-L4 and the genome-transplanted strain at 25°C. Individual values and standard deviations are shown as filled circles and error bars, respectively. *P* value of Student’s *t*-test is upper of graph.

## Discussion

### Mechanisms of low-temperature adaptation in the minimal cell

Here, we performed an adaptive laboratory evolution and obtain evolved JCVI-syn3B strains capable of growth at 25 °C, a temperature at which the original strain could not grow. This study demonstrated that minimal cells retain the capacity for rapid adaptation, highlighting their inherent evolvability despite a reduced genome and limited regulatory systems. To analyze detailed biological characteristics of the evolved strains, genome sequencing, RNA-seq, methylation analysis, and proteomic profiling were performed. Among these, the proteomic changes were considered to provide key insights into the mechanisms of low-temperature adaptation. Three major mechanisms of cold adaptation were inferred from the RNA-seq and proteomic data (Fig. 6).

**Figure 6.**
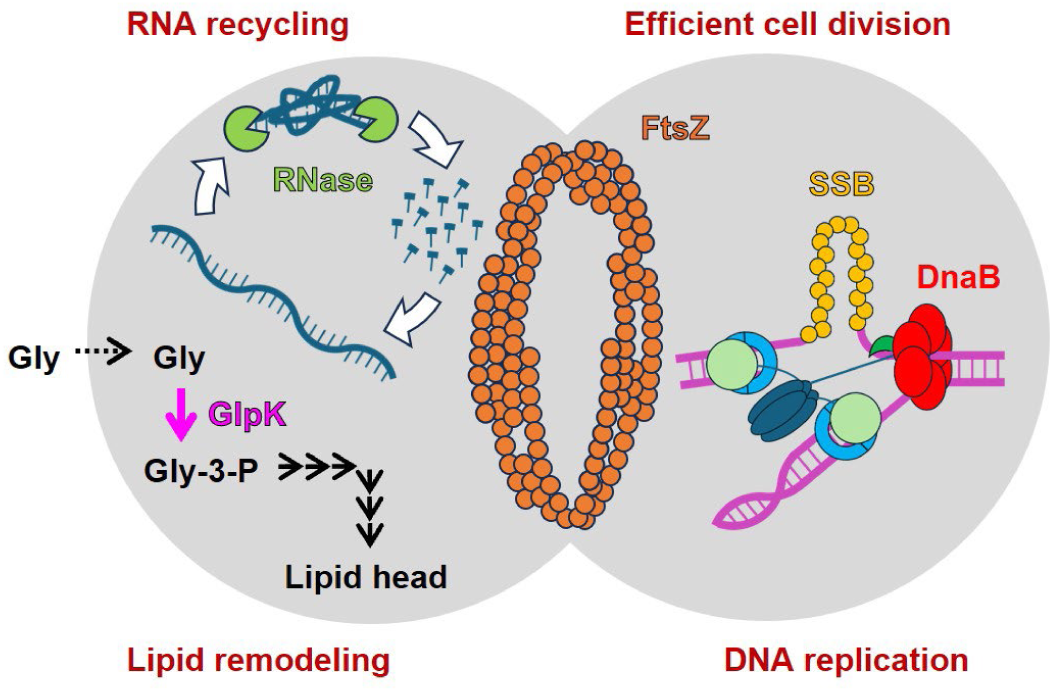
Schematic representation of the proposed mechanisms of cold adaptation. Four major mechanisms underlying cold adaptation were proposed in this study: (1) enhanced mRNA unwinding and recycling mediated by RNases, (2) maintenance of single-stranded DNA during replication, (3) lipid synthesis through glycerol metabolism, and (4) efficient cell division due to elevated expression levels of FtsZ and SepF. Several other mechanisms were also proposed.

### 1. mRNA unwinding and recycling

At low temperature, transcribed mRNAs are prone to forming secondary structures and double-stranded helices, either within themselves or through interactions with other mRNAs, resulting in the production of abnormal mRNAs. In this study, RNase levels were markedly elevated, with an average abundance of 240% in the cold-adapted strains (Extended Data Table 5). This substantial increase in RNases was presumably intended to actively degrade and recycle abnormal mRNAs. These findings were consistent with previous reports of the cold shock response in *E. coli* showing that the expression of CspA, a nucleotide-binding protein that binds to mRNA, and RNase R, which prevents the accumulation of abnormally folded mRNA ^7^.

### 2. Maintenance of single-stranded DNA

Low temperature can excessively stabilize the pairing of double-stranded DNA, as well as RNA. During DNA replication, unwinding and maintaining single-stranded DNA are essential ^21^. Our results showed that the protein levels of DNA helicase (DnaB) and single-stranded DNA-binding protein (SSB) were significantly increased in cold-adapted strains (Extended Data Table 6). These proteins likely facilitate DNA unwinding and stabilize single-stranded regions, thereby enabling efficient DNA replication. To our knowledge, this mechanism has not been previously reported in the context of cold adaptation.

### 3. Enhanced lipid metabolism

Transcriptomic and proteomic analyses revealed a substantial increase in *glpK*, which converts glycerol to glycerol-3-phosphate (Extended Data Table 3 and 6). Transcription levels of downstream pathways responsible for the synthesis of phosphatidic acid, phosphatidylglycerol, and cardiolipin were also generally increased (Supplementary Information Table 1). However, several enzymes in these pathways were not detected in the proteomic analysis, likely because they are membrane-associated proteins. These results suggest that enhanced lipid metabolism contributes to cold adaptation.

### 4. Cell division

Our results showed that most proteins involved in cell division, such as FtsZ, SepF, and FtsA, were upregulated (Extended Data Table 7). The assembly of FtsZ into the Z-ring and its subsequent contraction are essential for effective cell division ^22^. A reduction in the intrinsic GTPase activity of FtsZ proteins may impair Z-ring contraction. The observed upregulation of FtsZ at low temperature probably compensate for reduced GTPase activity by increasing the number of FtsZ molecule. Additionally, the elevated level of SepF, which anchors FtsZ to the cell membrane, may also play a critical role in maintaining the reinforced rings. Collectively, these higher protein levels would be important for efficient cell division under low-temperature conditions. To our knowledge, this mechanism has not been previously reported in the context of cold adaptation.

### 5. Other mechanisms

Several additional mechanisms were also suggested. For example, the protein levels of RbfA, which is required for 16S rRNA processing and cell growth at low temperature in *E. coli* ^23^, and RpsO, which encodes the 30S ribosomal protein S15 that functions as a scaffold for ribosome assembly ^24^, were markedly increased (Extended Data Fig. 3). These findings were consistent with previous reports showing upregulation of the *metY–rpsO* operon, which includes *rbfA* and *rpsO*, in *E. coli* under low-temperature conditions ^25,26^. The efficient rRNA processing and ribosome assembly would be important for cold adaptation in both JCVI-syn3B and *E. coli*. Elevated protein levels of tRNA, mRNA, and rRNA biogenesis pathways were also observed, indicating an increased demand for transcription and translation at low temperature. Moreover, the levels of F_o_ C-subunit, forming a membrane embedded rotational ring for establishment of membrane potential, and lipoproteins were substantially increased, implying potential roles for these proteins in cold adaptation.

These results suggested multiple mechanisms underlying cold adaptation in the minimal cell, including several potentially novel mechanisms. Abnormal RNA and DNA secondary structures may impose an energetic burden due to the need for degradation and recycling. However, over a long evolutionary process, the accumulation of mutations that reduce the tendency of mRNA or genomes to form abnormal secondary structures could minimize such costs. Natural psychrotrophic bacteria may have reached such an optimized, energy-efficient state, representing the fittest state. In contrast, the minimal cell adapted to low temperature while retaining this burden, representing an intermediate stage toward optimal fitness. This study provides an experimental snapshot of this evolutionary trajectory.

## Conclusion

This study demonstrates that even a minimal cell can evolve robustly under thermal constraint and provides insight into the core principles of cold adaptation. Despite a reduced genome and limited regulatory systems, minimal cells retain high evolvability and can rapidly adapt to new environments, enabling the identification of fundamental mechanisms of adaptation. The ability to distinguish genetic and non-genetic contributions through genome transplantation further enhances their experimental utility. In addition, low-temperature-adapted minimal cells provide a versatile platform for gene expression and functional analyses at near-ambient temperatures (Supplementary Discussion). Collectively, these findings establish minimal cells as a powerful platform for evolutionary and synthetic biology. Future studies under diverse environmental conditions will further expand their utility and reveal general principles of cellular adaptation.

---

**Extended Data Figure 1.**
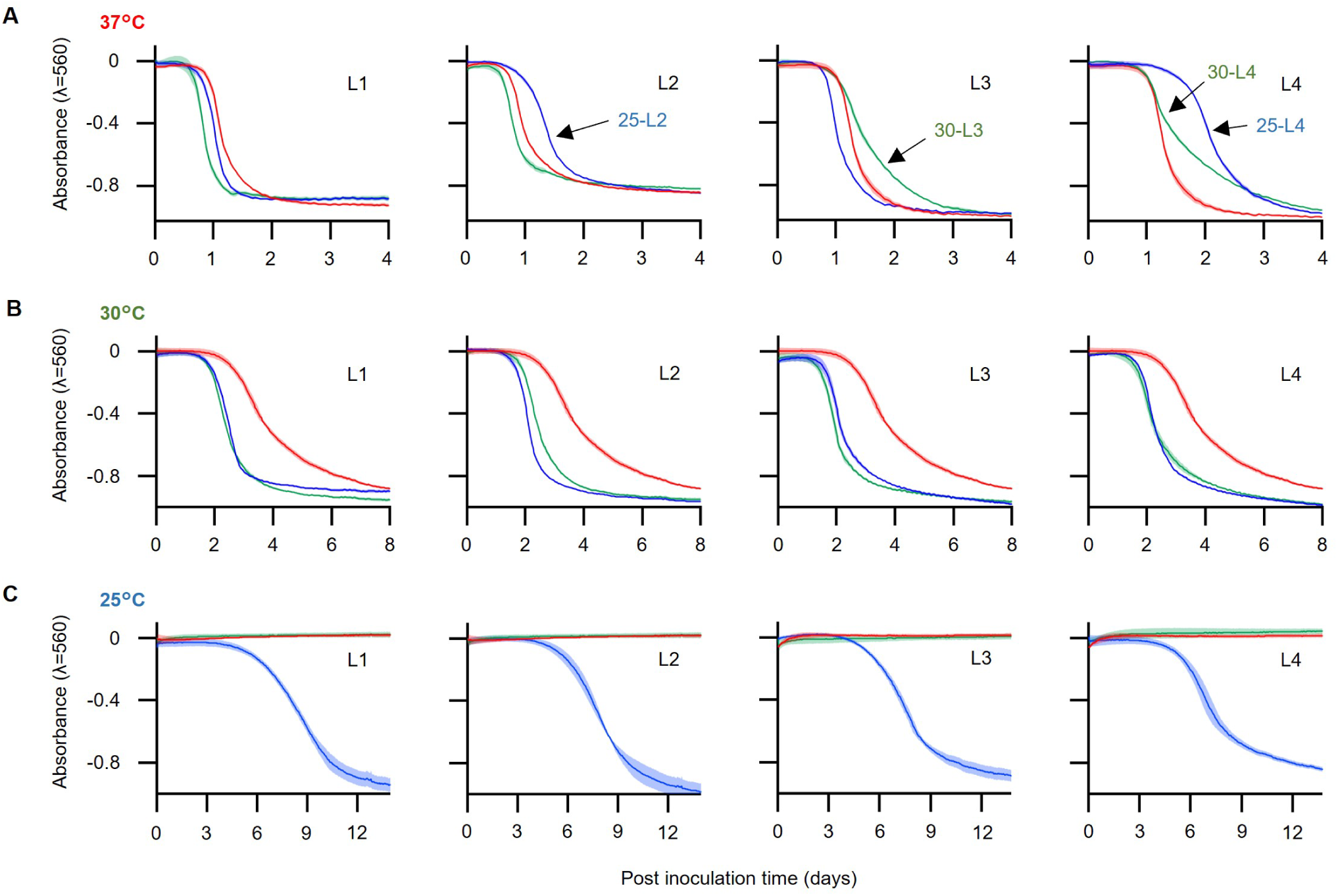
Growth curves of individual adapted strains. The original strain and the strains adapted to 30°C and 25°C are shown as red, green, and blue lines, respectively, during cultivation at 37°C (A), 30°C (B), and 25°C (C). The results of the four lineages were shown separately. The values are averages of three independent cultures, and the standard deviations are indicated by light-coloured lines. Four adapted lineages that exhibited a trade-off at 37°C are shown in panel A.

**Extended Data Figure 2.**
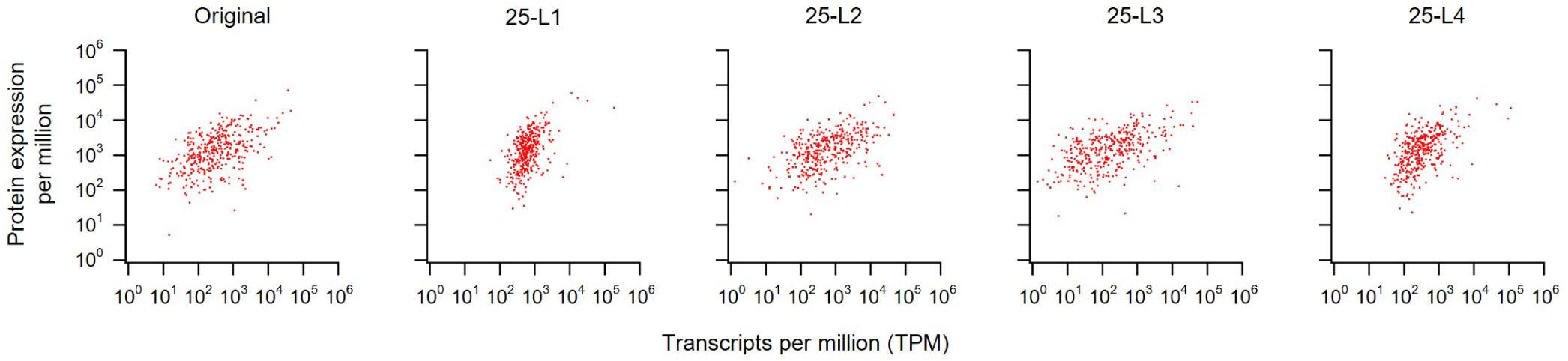
Correlation between RNA-seq and proteomics data. Expression levels from RNA-seq and proteomics datasets from the original and four 25°C-adapted strains were compared. The x-axis represents transcripts per million (TPM), and the y-axis indicates protein expression per million.

**Extended Data Figure 3.**
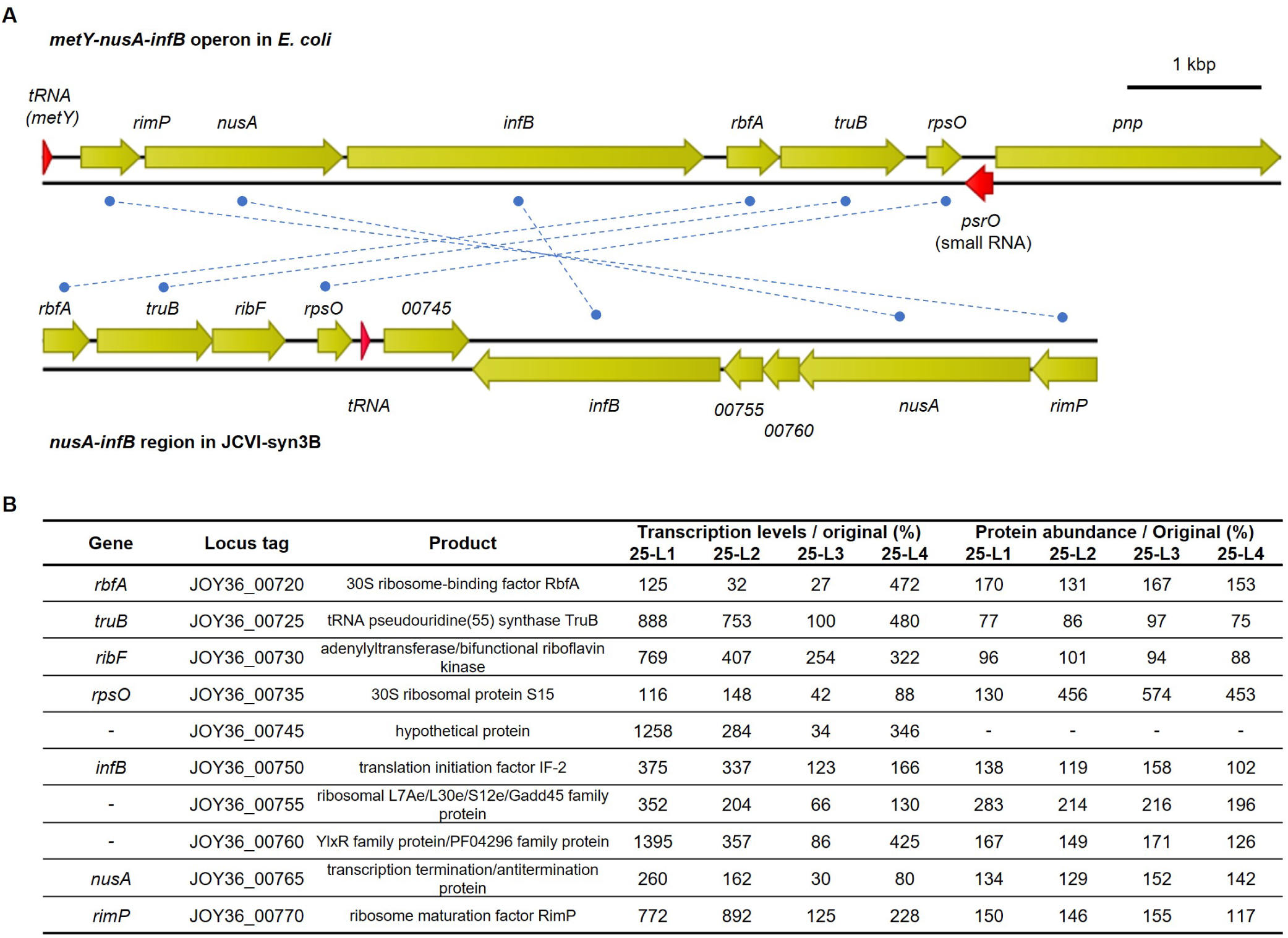
(A) Gene organization of the operon known to be upregulated during cold adaptation in *E. coli* and the corresponding homologous genomic region in JCVI-syn3B. Genes annotated with the same names are connected by blue dotted lines. (B) RNA-seq and proteomics data for genes encoded in this region in JCVI-syn3B. Most genes in this region were upregulated after cold adaptation in JCVI-syn3B.

**Extended Data Table 1.**
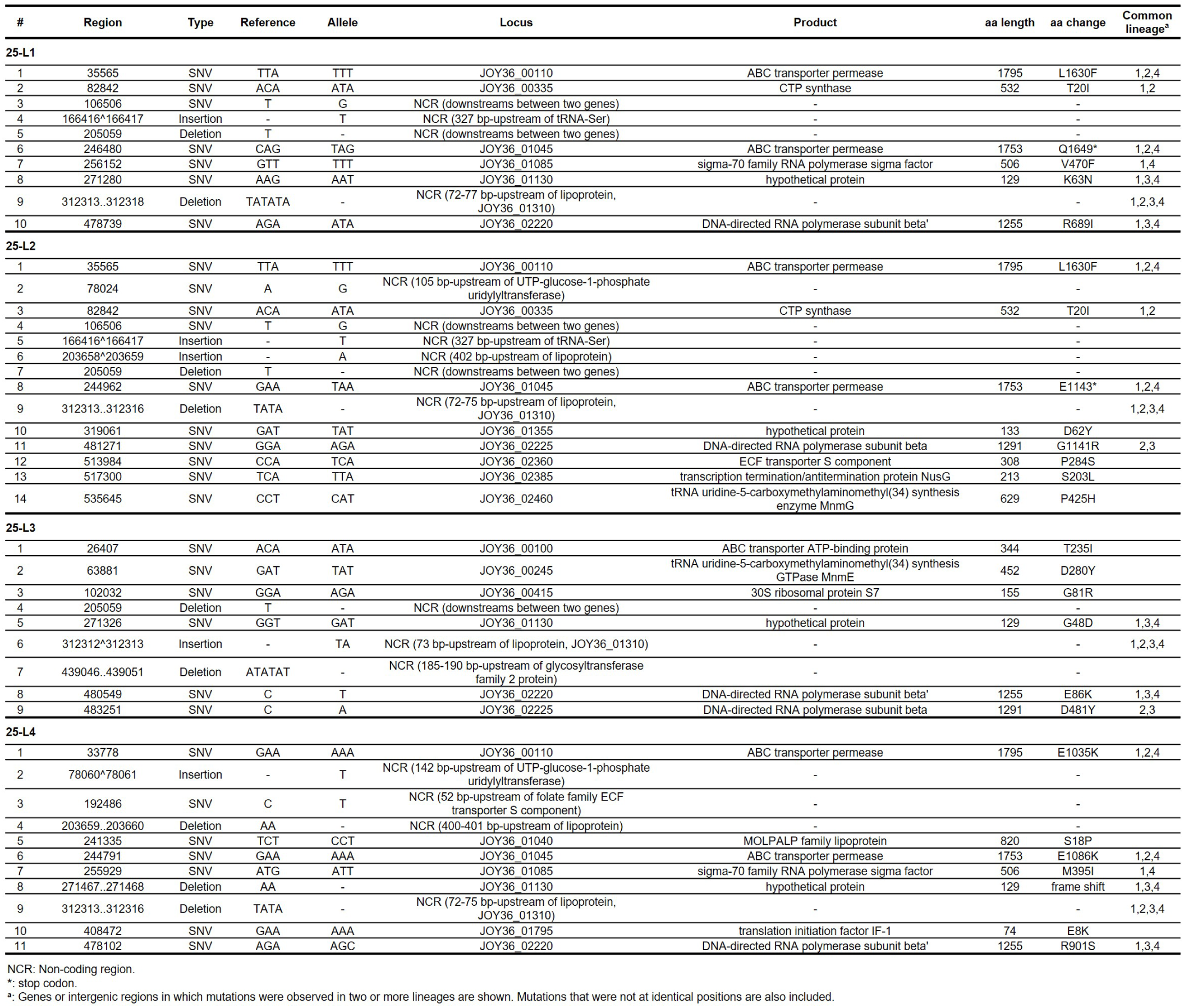
The genomic mutations detected in 25 ℃-adaped strains.

**Extended Data Table 2.** Transcription changes in the 25 ℃-adaped strains.

| Functional categories | Categorized gene count | Transcription levels / Total (%) |  |  |  |  |  | Transcription levels / original strain (%) |  |  |  |  |
| --- | --- | --- | --- | --- | --- | --- | --- | --- | --- | --- | --- | --- |
|  |  | Original | 25-L1 | 25-L2 | 25-L3 | 25-L4 | average | 25-L1 | 25-L2 | 25-L3 | 25-L4 | average |
| Metabolism |  |  |  |  |  |  |  |  |  |  |  |  |
| Glycolysis core | 10 | 1.69 | 1.54 | 2.52 | 1.53 | 1.72 | 1.83 | 91 | 149 | 90 | 102 | 108 |
| Pyruvate metabolism | 5 | 4.10 | 3.05 | 4.22 | 5.12 | 5.71 | 4.52 | 74 | 103 | 125 | 139 | 110 |
| Glycolysis branch | 5 | 0.53 | 0.77 | 0.58 | 0.15 | 0.35 | 0.46 | 146 | 110 | 29 | 66 | 88 |
| Pentose phosphate | 6 | 1.01 | 0.57 | 0.46 | 0.60 | 0.37 | 0.50 | 56 | 45 | 59 | 37 | 49 |
| Lipid metabolism | 20 | 2.02 | 4.13 | 3.48 | 1.77 | 11.72 | 5.28 | 204 | 172 | 88 | 581 | 261 |
| Nucleotide metabolism | 19 | 5.89 | 1.53 | 4.32 | 2.95 | 0.80 | 2.40 | 26 | 73 | 50 | 14 | 41 |
| Amino acid metabolism | 9 | 0.26 | 0.46 | 0.23 | 0.15 | 0.29 | 0.28 | 175 | 88 | 57 | 109 | 107 |
| Cofactor metabolism | 10 | 0.15 | 0.62 | 0.34 | 0.09 | 0.33 | 0.35 | 409 | 225 | 61 | 215 | 228 |
| Macromolecule metabolism | 3 | 0.81 | 0.33 | 1.05 | 0.66 | 0.38 | 0.61 | 41 | 130 | 82 | 47 | 75 |
| Environmental Information Processing |  |  |  |  |  |  |  |  |  |  |  |  |
| PTS | 4 | 2.21 | 18.90 | 5.62 | 5.13 | 10.11 | 9.94 | 856 | 255 | 233 | 458 | 450 |
| Cofactor | 15 | 0.37 | 1.75 | 1.81 | 0.49 | 1.60 | 1.41 | 472 | 488 | 133 | 430 | 381 |
| Macromolecule | 12 | 0.41 | 0.79 | 0.54 | 0.15 | 0.28 | 0.44 | 193 | 131 | 36 | 69 | 107 |
| Ion (Pi, Ca, Mg, K) | 7 | 0.25 | 0.34 | 0.23 | 0.10 | 0.19 | 0.22 | 138 | 92 | 41 | 77 | 87 |
| F-type ATPase | 8 | 0.33 | 0.64 | 1.85 | 0.41 | 0.54 | 0.86 | 197 | 565 | 127 | 164 | 263 |
| Genetic Information Processing |  |  |  |  |  |  |  |  |  |  |  |  |
| DNA replication | 17 | 0.39 | 0.97 | 0.93 | 0.25 | 0.52 | 0.67 | 245 | 236 | 65 | 132 | 169 |
| DNA repair | 11 | 0.21 | 0.49 | 0.60 | 0.11 | 0.20 | 0.35 | 240 | 295 | 55 | 99 | 172 |
| tRNA biogenesis | 19 | 31.94 | 34.65 | 8.66 | 29.91 | 33.42 | 26.66 | 108 | 27 | 94 | 105 | 83 |
| rRNA biogenesis | 18 | 0.41 | 0.81 | 0.84 | 0.32 | 0.41 | 0.59 | 200 | 206 | 78 | 102 | 146 |
| Transcription | 12 | 1.72 | 0.80 | 2.05 | 0.65 | 0.49 | 1.00 | 46 | 119 | 38 | 28 | 58 |
| RNA degradation | 6 | 0.27 | 0.74 | 2.24 | 0.41 | 0.61 | 1.00 | 272 | 817 | 148 | 223 | 365 |
| Translation cycle | 14 | 18.02 | 10.05 | 9.98 | 23.19 | 17.86 | 15.27 | 56 | 55 | 129 | 99 | 85 |
| Ribosomal protein | 54 | 13.52 | 4.13 | 27.90 | 12.59 | 3.54 | 12.04 | 31 | 206 | 93 | 26 | 89 |
| tRNA-ligase | 20 | 0.74 | 1.17 | 1.04 | 0.28 | 0.64 | 0.78 | 159 | 141 | 38 | 86 | 106 |
| Chaperone | 3 | 0.30 | 0.20 | 0.25 | 0.07 | 0.18 | 0.18 | 68 | 83 | 24 | 58 | 58 |
| Protein export | 9 | 0.55 | 0.76 | 2.09 | 0.26 | 0.32 | 0.86 | 138 | 380 | 47 | 59 | 156 |
| Cellular Processes |  |  |  |  |  |  |  |  |  |  |  |  |
| Cell division | 5 | 1.95 | 1.45 | 1.94 | 5.31 | 1.61 | 2.58 | 74 | 100 | 273 | 83 | 132 |
| Chromosome condensation and segregation | 6 | 0.24 | 0.23 | 0.08 | 0.03 | 0.11 | 0.11 | 98 | 33 | 13 | 47 | 48 |
| Others |  |  |  |  |  |  |  |  |  |  |  |  |
| Lipoprotein | 13 | 0.99 | 0.89 | 1.43 | 0.73 | 1.02 | 1.02 | 91 | 146 | 74 | 103 | 103 |
| Redox | 3 | 1.18 | 0.34 | 0.69 | 0.62 | 0.28 | 0.48 | 29 | 58 | 52 | 24 | 41 |
| Hypothetical protein | 52 | 2.45 | 2.47 | 3.35 | 2.30 | 1.12 | 2.31 | 101 | 136 | 94 | 46 | 94 |
| Unclassified | 69 | 5.09 | 4.40 | 8.69 | 3.64 | 3.29 | 5.00 | 86 | 171 | 71 | 65 | 98 |

**Extended Data Table 3.**
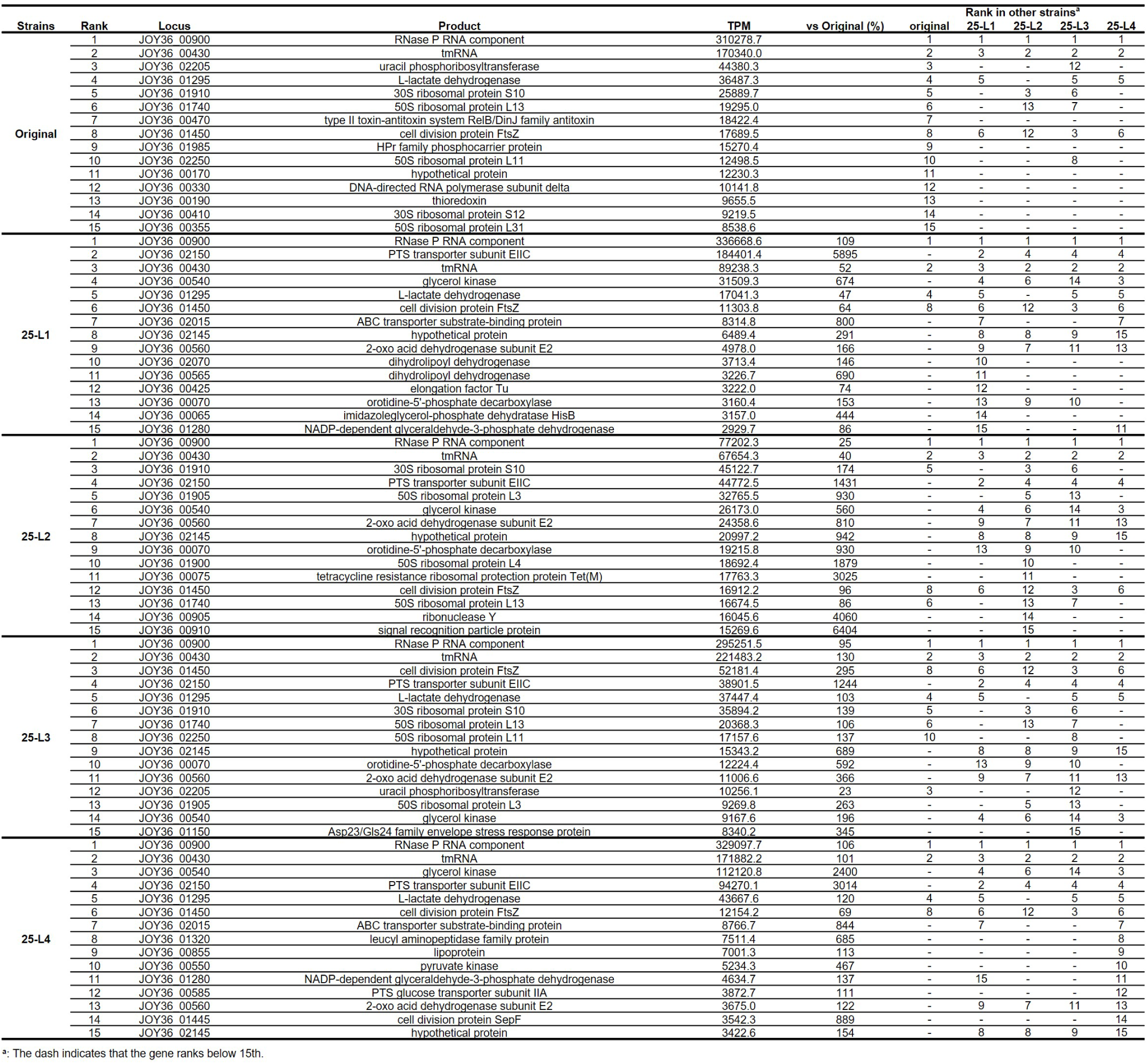
Genes with high transcription levels in each strain.

**Extended Data Table 4.** Total number of methylated sites in the genomes of the original and the 25 ℃-adaped strains.

| Methylation type | Original #1<sup>a</sup> | Original #2<sup>a</sup> | 25-L1 #1<sup>a</sup> | 25-L1 #2<sup>a</sup> | 25-L4 #1<sup>a</sup> | 25-L4 #2<sup>a</sup> |
| --- | --- | --- | --- | --- | --- | --- |
| m6A | 593 | 588 | 586 | 560 | 574 | 573 |
| m4C | 5273 | 5285 | 5179 | 4902 | 5139 | 5046 |
| modified base<sup>b</sup> | 23884 | 24242 | 23565 | 22232 | 23238 | 23064 |

<sup>a</sup>. Whole genome methylation analysis were performed in duplicate.
<sup>b</sup>. Modification not recognized by the software.

**Extended Data Table 5.** Protein expression levels in the 25 ℃-adaped strains.

| Functional categories | Categorized protein count | Original | Expression levels / Total (%) |  |  |  | Expression levels / original strain (%) |  |  |  | average |
| --- | --- | --- | --- | --- | --- | --- | --- | --- | --- | --- | --- |
|  |  |  | 25-L1 | 25-L2 | 25-L3 | 25-L4 | 25-L1 | 25-L2 | 25-L3 | 25-L4 |  |
| Metabolism |  |  |  |  |  |  |  |  |  |  |  |
| Glycolysis core | 10 | 6.02 | 7.11 | 5.56 | 6.40 | 6.19 | 118 | 92 | 106 | 103 | 105 |
| Pyruvate metabolism | 5 | 8.81 | 5.73 | 4.35 | 5.00 | 4.37 | 65 | 49 | 57 | 50 | 55 |
| Glycolysis branch | 4 | 0.47 | 0.41 | 0.46 | 0.44 | 0.41 | 87 | 98 | 94 | 88 | 92 |
| Pentose phosphate | 6 | 1.43 | 1.16 | 1.20 | 1.23 | 1.19 | 81 | 84 | 86 | 83 | 84 |
| Lipid metabolism | 14 | 2.76 | 5.34 | 5.41 | 3.54 | 4.30 | 193 | 196 | 128 | 156 | 168 |
| Nucleotide metabolism | 18 | 5.23 | 3.89 | 3.99 | 5.51 | 6.33 | 74 | 76 | 105 | 121 | 94 |
| Amino acid metabolism | 8 | 2.04 | 1.54 | 1.91 | 2.07 | 2.04 | 76 | 94 | 101 | 100 | 93 |
| Cofactor metabolism | 8 | 0.76 | 0.62 | 0.62 | 0.63 | 0.54 | 81 | 83 | 83 | 71 | 79 |
| Macromolecule metabolism | 3 | 0.60 | 0.60 | 0.61 | 0.58 | 0.62 | 101 | 102 | 97 | 103 | 101 |
| Environmental Information Processing |  |  |  |  |  |  |  |  |  |  |  |
| PTS | 4 | 2.86 | 3.36 | 2.56 | 1.69 | 2.11 | 117 | 90 | 59 | 74 | 85 |
| Cofactor | 7 | 1.29 | 1.33 | 1.91 | 2.01 | 2.35 | 103 | 149 | 156 | 183 | 148 |
| Macromolecule | 7 | 0.52 | 0.65 | 0.89 | 0.76 | 0.92 | 125 | 173 | 148 | 178 | 156 |
| Ion (Pi, Ca, Mg, K) | 5 | 0.49 | 0.61 | 0.72 | 0.71 | 0.75 | 124 | 148 | 145 | 154 | 143 |
| F-type ATPase | 7 | 0.54 | 0.63 | 0.71 | 0.58 | 0.68 | 116 | 130 | 107 | 126 | 120 |
| Genetic Information Processing |  |  |  |  |  |  |  |  |  |  |  |
| DNA replication | 15 | 2.61 | 2.17 | 3.18 | 3.48 | 3.90 | 83 | 122 | 133 | 150 | 122 |
| DNA repair | 8 | 0.82 | 0.69 | 0.65 | 0.69 | 0.64 | 84 | 80 | 84 | 78 | 82 |
| tRNA biogenesis | 17 | 1.38 | 1.24 | 1.76 | 1.83 | 1.89 | 90 | 128 | 133 | 137 | 122 |
| rRNA biogenesis | 17 | 1.32 | 1.87 | 2.06 | 1.95 | 1.81 | 141 | 155 | 147 | 136 | 145 |
| Transcription | 11 | 3.24 | 3.69 | 3.81 | 3.90 | 3.69 | 114 | 118 | 121 | 114 | 116 |
| RNA degradation | 6 | 1.27 | 2.64 | 2.92 | 3.17 | 3.42 | 209 | 230 | 250 | 270 | 240 |
| Translation cycle | 13 | 8.10 | 6.64 | 6.52 | 6.14 | 5.21 | 82 | 81 | 76 | 64 | 76 |
| Ribosomal protein | 47 | 24.47 | 20.02 | 21.42 | 22.23 | 19.50 | 82 | 88 | 91 | 80 | 85 |
| tRNA-ligase | 20 | 3.49 | 3.52 | 3.81 | 4.01 | 4.03 | 101 | 109 | 115 | 115 | 110 |
| Chaperone | 3 | 1.60 | 0.92 | 1.45 | 1.52 | 1.83 | 57 | 91 | 95 | 114 | 89 |
| Protein export | 7 | 1.00 | 0.97 | 0.99 | 0.94 | 1.04 | 97 | 99 | 94 | 104 | 98 |
| Cellular Processes |  |  |  |  |  |  |  |  |  |  |  |
| Cell division | 4 | 1.79 | 6.87 | 5.65 | 4.24 | 5.30 | 383 | 315 | 237 | 295 | 307 |
| Chromosome condensation and segregation | 5 | 0.49 | 0.50 | 0.52 | 0.47 | 0.43 | 102 | 105 | 97 | 89 | 98 |
| Others |  |  |  |  |  |  |  |  |  |  |  |
| Lipoprotein | 8 | 1.63 | 2.52 | 2.03 | 1.97 | 2.01 | 154 | 124 | 121 | 123 | 131 |
| Redox | 3 | 2.43 | 1.39 | 1.25 | 1.50 | 1.33 | 57 | 51 | 62 | 55 | 56 |
| Hypothetical protein | 27 | 1.79 | 2.31 | 2.27 | 2.09 | 2.31 | 129 | 127 | 117 | 129 | 125 |
| Unclassified | 54 | 8.76 | 9.06 | 8.82 | 8.69 | 8.88 | 103 | 101 | 99 | 101 | 101 |

**Extended Data Table 6.**
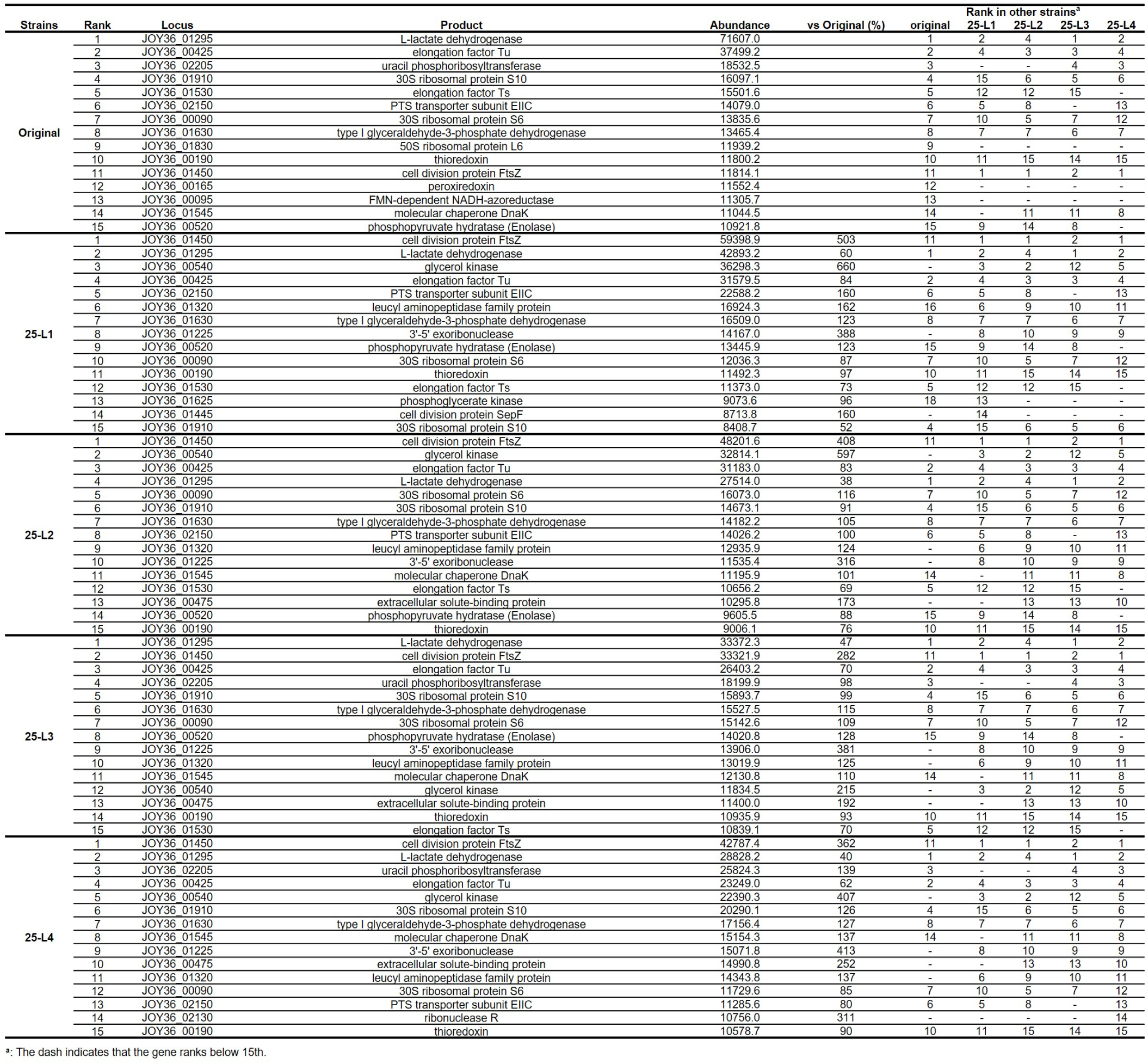
Protein with high abundance in each lineage.

**Extended Data Table 7.** Protein with substantially altered expression after cold adaptation at 25 ℃.

| Protein | Locus tag | Product | Protein abundance / Original (%) |  |  |  |  |
| --- | --- | --- | --- | --- | --- | --- | --- |
|  |  |  | 25-L1 | 25-L2 | 25-L3 | 25-L4 | average |
| Cell division |  |  |  |  |  |  |  |
| FtsZ | JOY36_01450 | cell division protein FtsZ | 503 | 408 | 282 | 362 | 389 |
| SepF | JOY36_01445 | cell division protein SepF | 160 | 140 | 151 | 173 | 156 |
| FtsA | JOY36_01455 | cell division FtsA domain-containing protein | 136 | 96 | 126 | 125 | 121 |
| DivIVA | JOY36_00895 | DivIVA domain-containing protein | 74 | 86 | 133 | 103 | 99 |
| RNA degradation |  |  |  |  |  |  |  |
| YhaM | JOY36_01225 | 3'-5' exoribonuclease | 388 | 316 | 381 | 413 | 375 |
| Rny | JOY36_00905 | ribonuclease Y | 123 | 141 | 108 | 115 | 122 |
| Rnr | JOY36_02130 | ribonuclease R | 75 | 239 | 297 | 311 | 230 |
| RnjB | JOY36_00635 | ribonuclease J | 217 | 195 | 158 | 188 | 190 |
| RnjA | JOY36_01600 | ribonuclease J | 135 | 155 | 114 | 113 | 129 |
| Lipid metabolism |  |  |  |  |  |  |  |
| GlpK | JOY36_00540 | glycerol kinase | 660 | 597 | 215 | 407 | 470 |
| PlsC | JOY36_01405 | 1-acyl-sn-glycerol-3-phosphate acyltransferase | 274 | 265 | 207 | 217 | 241 |
| - | JOY36_00300 | glycosyltransferase family 2 protein | 97 | 143 | 148 | 84 | 118 |
| - | JOY36_02000 | glycosyltransferase family 2 protein | 161 | 109 | 110 | 118 | 125 |
| GalU | JOY36_00305 | UTP--glucose-1-phosphate uridylyltransferase | 155 | 71 | 260 | 372 | 215 |
| DNA replication |  |  |  |  |  |  |  |
| DnaB | JOY36_01640 | replicative DNA helicase | 93 | 306 | 509 | 855 | 441 |
| Ssb | JOY36_00085 | single-stranded DNA-binding protein | 177 | 302 | 316 | 189 | 246 |
| DNA damage repair |  |  |  |  |  |  |  |
| UvrB | JOY36_02310 | excinuclease ABC subunit UvrB | 219 | 234 | 258 | 182 | 223 |

## METHODS

### Bacterial strain and adaptive laboratory evolution experiments

The synthetic bacterium JCVI-syn3B (GenBank, CP069345.1) was cultured in SP4 medium containing phenol red at 37°C 1. The JCVI-syn3B is a derivative strain with 20 additional genes compared to JCVI-syn3.0 2,3. This strain showed relatively normal cell division and morphology, and growth speed, making it useful for research 2,3. Since JCVI-syn3B is derived from M. mycoides known as a parasitic bacterium of cattle, its optimal growth temperature, 37°C, is around the body temperature of the host animal, 38‒39°C 4. The color of the culture medium changes from red to yellow as cell growth proceeds, due to the production of acidic metabolites.

For the adaptive laboratory evolution experiments, the original JCVI-syn3B strain was first cultured at 37°C in SP4 medium. After the culture color changed from red to yellowish orange, 5 μL of culture was transferred to 500 μL of fresh SP4 medium and incubated at 30°C (passage 1 at 30°C). This process was repeated every time when the culture changed color, up to passage 20, in four parallel lineages. After 20 passages at 30°C, 5 μL of culture was transferred to 500 μL of fresh SP4 medium and incubated at 25°C (passage 1 at 25°C). This process was also repeated up to passage 20 in four parallel lineages.

### Growth measurement

The cultured bacterial lines were monitored by the absorbance at 560 nm wavelength that is strongly correlated with color change of phenol red 1. To each well of a 96-well plate, 10 ml cell culture containing around 5.0×105 cells was inoculated to 190 μL fresh SP4 medium, whose top side was sealed with LightCycler 480 Sealing Foil (Nippon Genetics, Tokyo, Japan). The plate was set to a microplate reader Absorbance 96 (Byonoy, Hamburg, Germany), and each culture was monitored by the absorbance at 560 nm for 6 days every hour. The absorbance values of cultures were calibrated by subtracting the absorbance value of plain medium as control. The doubling time was calculated by linear fitting of the early exponential growth phase, based on the time required for the absorbance value to change from −0.2 to −0.4. To investigate the growth rates of the evolved strains at 25°C, 30°C and 37°C, frozen stocks with known colony-forming units (CFUs) were used to ensure uniform inoculum size across all experiments. The frozen stocks were thawed and diluted with SP4 medium to 2.5×105 cells in 10 μL, which were then inoculated to 190 μL of SP4 medium and cultured at 37°C, 30°C or 25°C. For growth comparison between the evolved strains and the original strain, 10 μL of cell culture containing 2.5×105 cells were inoculate, and for the genome-transplanted, evolved, and original strains, 10 μL of cell culture containing 5.0×105 cells were inoculated to 190 μl of SP4 medium, and subjected to the measurements, respectively.

### Genome sequencing

The passaged strains were cultured on SP4 agar plates, and single colonies were picked and grown in SP4 liquid medium. The cultured cells were collected by centrifugation at 9000 g for 8 min at 22℃, washed with S/T buffer (0.5 M sucrose, 10 mM Tris-HCl; pH 6.5), then subjected to DNA extraction using NucleoSpin Tissue kit (Macherey-Nagel, Duren, Germany) according to the manufacturer’s instructions and sequencing using DNBSEQ platform (MGI Tech, Shenzhen, China). The obtained reads were mapped to the JCVI-syn3B genome (GenBank: CP069345), and mutations were identified using CLC Genomics Workbench version 21 software (Qiagen, Hilden, Germany).

### Protein profiling and mass spectrometry

For SDS-PAGE analysis, the cultured cells were collected, washed with S/T buffer, lysed by BactYeast Lysis buffer (ATTO, Tokyo, Japan), mixed with EzApply (ATTO), and heated at 95°C for 5 min. The samples were loaded onto a 5%‒20% gradient polyacrylamide gel and electrophoresed at 250 V and 20 mA for 80 min. The gel was stained with Coomassie Brilliant Blue solution and imaged using a scanner (GT-X830; Epson, Tokyo, Japan). The protein bands in the gel were excised, and subjected to protein identification as described previously 5. In this study, the digested and extracted proteins from the gel were analyzed using a nanoLC-MS system composed of a nanoflow ultra-high-performance liquid chromatography (Acquity UPLC M-Class, Waters) and a quadrupole time-of-flight mass spectrometry (Xevo G2-XS qTOF, Waters) equipped with an electrospray ionization source. The digested peptides were separated using a nanoLC column (nanoEase HSS T3, 75 µm×150 mm, Waters) under a water-acetonitrile gradient at a flow rate of 350 nL/min. Mass spectra were acquired in positive MSe modeIn re. Protein identification was performed using ProteinLynx Global Server (Waters) with reference to JCVI-syn3B protein sequences.

For proteome analysis, an aliquot of cultured medium estimated to contain 50 µg protein was mixed with four volumes of acetone, and the precipitated protein pellet was subjected to in-solution digestion as described previously 6. The same nanoLC-MS system and MSe acquisition mode as above were used. The obtained mass data were analyzed with Progenesis QIP software (Waters). Four technical replicates were performed for each evolved strain.

### RNA-seq analysis

The cultured cells were collected and washed with TE buffer (10 mM Tris-HCl, 1 mM EDTA, pH8.0). The total RNA samples was extracted using NucleoSpin RNA (Macherey-Nagel) by the standard protocol and their concentrations were measured using Qubit fluorometer with RNA assay kit (invitrogen). Ribosomal RNAs were depleted by Ribo-off rRNA Depletion Kit V2 (Bacteria) (Vazyme, Nanjing, China). The libraries were constructed by MGIEasy Fast RNA Library Prep Set (MGI Tech) and circularized by MGIEasy Dual Barcode Circularization Kit (MGI Tech). DNA nanoballs were constructed by DNB Rapid Make Reagent Kit (MGI Tech) and High-throughput Pair-End Sequencing Primer Kit (App-D) (MGI Tech) and sequenced using DNBSEQ-T7 (MGI Tech). Obtained reads were analyzed by CLC Genomics Workbench 25 software (Qiagen).

### Transformation and genome transplantation

For introduction of a puromycin-resistant gene into the bacterial genome, Cre/lox-mediated transformation method was used as previously described 1. The bacterial cells were collected by centrifugation at 9,000 g for 8 min at 22℃, washed with S/T buffer, collected and suspended 0.1 M CaCl2 solution and incubated for 30 min on ice. The competent cells were mixed with 10-100 ng pSK020 plasmids and 70% polyethylene glycol. The transformants were screened on SP4 agar plates containing 3 μg/mL puromycin. Genomic DNA extraction and genome transplantation were performed using a previously described method 7. The bacterial cells were collected from 20 ml of well-grown SP4 medium by centrifugation, washed with 10 mL of S/T buffer, collected by centrifugation, and suspended in 500 μL of S/T buffer. The cell suspension was mixed with 500 μL of 2% low-melting-point agarose solution, of which 90 μL was applied to each plug mold (Bio-Rad, California, U.S.). After solidification at 4°C, the plugs were treated with 5 mL of Proteinase K reaction buffer (0.1 M EDTA [pH 8.0], 1% sodium dodecyl sulfate, 20 mg/mL Proteinase K) at 50°C for three days with reaction buffer replacement per day. The treated plugs were washed with 20 mL of Tris-EDTA buffer (20 mM Tris-HCl and 50 mM EDTA [pH 8.0]) for four hours under gentle agitation with buffer replacement per hour.

### DNA methylation analysis

Whole-genome methylation analysis was performed for the original, 25-L1, and 25-L4 strains. The cultured cells were collected, washed with S/T buffer, then subjected to DNA extraction using NucleoBond HMW DNA kit (Macherey-Nage) according to the manufacturer’s instructions. Sequencing was outsourced to Bioengineering Lab. Co., Ltd, Japan. DNA methylation sites were identified during genome sequencing with a PacBio sequencer 8,9. Duplicate data were obtained for each strain. Analyses were conducted for m4C and m6A, which are representative types of methylation in bacteria, as well as for other types of methylation sites. Mapping of methylation sites to each gene was carried out using Microsoft Excel.

## Data Availability

Raw reads for whole genome sequencing and RNA-seq were deposited in DDBJ as accession codes DRA021582‒DRA021586 and DRA021615, respectively.

## Author Contributions

SK and Mizutani M conceived the study. SK, RK and Mizutani M designed experimental plans. Mizutani M performed almost all experiments. Moriyama M performed proteomics analysis. Mizutani M and SK performed data analysis. SK, Mizutani M and TF wrote the paper. All authors edited the paper and approved the final version of the manuscript.

## Funding

This study was supported by the JST ERATO Grant Number JPMJER1902 to SK and TF, and the JSPS KAKENHI Grant 18H02433, 26710015, 26106004, 15KK0266 to SK, JP17H06388 to TF, 24K18102 to Mizutani M, AMED Grant Number JP23gm1610002 to SK, and Research funds from Yamagata Prefecture and Tsuruoka City. Mizutani M was supported by the JSPS Research Fellowships for Young Scientists (22KJ318 to Mizutani M).

## Acknowledgements

We thank Yoko Hagiwara for supporting passaged cultivations.

## Notes

### Competing Interest Statement

The authors have declared no competing interest.

